# *In-situ* Target Base Editing Combining with Biosensor-driven Strategy Reveals Critical Single Nucleotide Variants for Enhanced Recombinant Protein Secretion in *Pichia pastoris*

**DOI:** 10.64898/2026.04.09.717583

**Authors:** Yaomeng Yuan, Xihao Liao, Yujing Tang, Shuang Li, Kaho Hasegawa, Yi Wang, Xin-Hui Xing, Chong Zhang

## Abstract

The disparity between the production and demand of recombinant proteins (r-proteins) has significantly hindered their commercial viability. Leveraging genomic resources offers substantial promise in enhancing our comprehension of metabolic and regulatory networks, thus facilitating the development of highly productive protein cell factories. However, the considerable gap between high-throughput strategies for monitoring r-protein secretion and genome perturbation in *P. pastoris* continues to obstruct the systematic linkage of genotype and phenotype, thereby limiting the optimization of production. Here, we developed a novel strategy combining dual-base editor-mediated *in-situ* genome engineering with nanobody-regulated biosensor-assisted droplet sorting to enhance r-protein secretion (BINDER) in *P. pastoris*. We successfully employed BINDER to screen recombinant human serum albumin (rHSA) hyper-producers and identified two critical SNVs conferring up to a 1.78-fold improved secretion titer from 113,632 mutants, providing valuable insights into the secretion mechanism. Fed-batch cultivation of the engineered strain resulted in the highest reported rHSA titer, 23.43g/L, in *P. pastoris*, demonstrating its substantial potential for industrial applications. Given the high transferability of base editors and the novel biosensor’s independence from the properties of the target protein, the strategy developed here might be expanded to a variety of microbial species and r-proteins.

## 1. Introduction

Recombinant proteins (r-proteins) have revolutionized various industries, with their market size and impact expected to continue growing(Rettenbacher et al., 2022). The methylotrophic yeast *Komagataella phaffii*, also known as *Pichia pastoris*, is a promising host for the hyper-production of r-proteins due to its high cell-density cultivation, robust secretory capabilities, efficient post-translational modification processes, and cost-effective downstream purification(Karbalaei et al., 2020). To date, over 5,000 r-proteins have been successfully expressed in *P. pastoris*(Schwarzhans et al., 2017). Strategies including codon optimization, dosage optimization, signal peptide optimization, as well as modification of a handful of genes related to secretion have been utilized to enhance the r-protein production in *P. pastoris*, realizing significant improvement in the titer of target r-proteins(Che et al., 2020; Liao et al., 2020; Lin et al., 2013; Mellitzer et al., 2014; Vogl et al., 2016; Zahrl et al., 2019; Zahrl et al., 2022; Zhu et al., 2018). Despite these successes, most of the engineered strains are not commercially viable, particularly for the production of protein-based biological materials, bulk food proteins, and bulk pharmaceutical proteins(Miserez et al., 2023; Piazenski et al., 2024; Werten et al., 2019). The annual demand for these products ranges from hundreds to hundreds of thousands of tons(Moghaddassi et al., 2014; Yang et al., 2017), whereas the highest r-protein secretion reported for *P. pastoris* is only 17∼20 g/L(Mellitzer et al., 2014; Zhu et al., 2021), hindering their large-scale application. Enhancing r-protein production offers a viable pathway to mitigate costs(Edlund et al., 2018; Piazenski et al., 2024; Werten et al., 2019). However, due to the lack of biological knowledge and the complexity of expression and secretion mechanism, the application of traditional strategies in further improving r-protein secretion titer is limited. Recent genome-wide researches have revealed numerous unknown genetic loci and biological processes within the genome that affect r-protein secretion and expression(Huang et al., 2015; Huang et al., 2017; Ito et al., 2022). This suggests that fully exploiting genomic resources holds great potential in advancing our understanding of metabolic and regulatory networks, thereby paving the way for the construction of the r-protein cell factory with ultra-high productivity. However, limited progress has left a significant gap between high-throughput monitoring of r-protein secretion and genome perturbation, hindering the systematic association of genotype and phenotype in *P. pastoris*, thereby limiting production improvements.

CRISPR-Cas9-derived strategies enable the modification of gene expression levels or gene loss of function through indel formation, thereby facilitating the acquisition of biological knowledge and the optimization of traits at the single-gene level(Liao et al., 2021; Tafrishi et al., 2024). However, these methods often limit information gain to lower resolution, restricting our understanding of the roles of specific positions within a gene. *In-situ* evolution of target genes have been proved to enhance traits such as stress tolerance and production capacity in microbial cell factories, underscoring the pivotal role of single nucleotide variants (SNVs) in determining phenotypes(Deng et al., 2024; Garst et al., 2017; Wang et al., 2009). To apply similar strategies in *P. pastoris*, CRISPR-Cas system mediated recombineering strategies have been established and optimized(Cai et al., 2021; Wang et al., 2023; Zhang et al., 2021). Recent advances in this method have enabled efficient gene editing using donor DNA with homology arms less than 50 bp by introducing proteins responsible for recombination(Deng et al., 2023; Gao et al., 2022). However, these approaches often suffer from the complexity of strain and plasmid construction, high costs of synthesizing donor DNA pools, and limited transformation and recombination efficiency, impeding the generation of extensive mutant libraries in *P. pastoris.* A promising alternative is base editors (BEs), which enable the introduction of SNVs by directly modifying target bases rather than replacing DNA fragments. Since complex DNA libraries are created *in vivo*, mutants with extensive genomic diversity can be obtained through serial passaging, thus overcoming the limitations of CRISPR-Cas9 mediated recombineering. By fusing cytosine or adenine deaminases with a catalytically impaired Cas nuclease, cytosine base editors (CBEs) or adenine base editors (ABEs) have been created to produce C>T or A>G conversions(Anzalone et al., 2020). Combining both cytosine and adenine deaminases, dual-base editors (DBEs) hold greater potential for generating more variations(Grünewald et al., 2020; Hao et al., 2022; Li et al., 2020; Sakata et al., 2020; Shelake et al., 2023; Xie et al., 2020; Zhang et al., 2020a; Zhang et al., 2020b).

CRISPR-system-based approaches enable the generation of millions of mutants, necessitating the advancement of high-throughput secretory phenotypic assays for the isolation and identification of hyper-secretors. Fluorescent reporter systems that convert protein production into a detectable fluorescent signal have facilitated high-throughput screening mediated by fluorescence-activated cell sorting (FACS) or fluorescence-activated droplet sorting (FADS). For secreted proteins, specific substrates coupled with fluorescent signals have been applied to screen enzyme hyper-secretors in droplet microfluidics systems(Baret et al., 2009; Johansson et al., 2023; Najah et al., 2014; Sjostrom et al., 2014). However, these strategies rely on the unique properties of enzymes, limiting their applicability to a broader range of r-proteins. To address this issue, strategies have been proposed by fusing a specific tag to the r-protein, thereby enabling the excitation of fluorescence when extracellular probes are added. Representatives including split GFP sensors(Knapp et al., 2017), Quenchbodies(Abe et al., 2011) and FlAsH-tetracysteine assay(Haitjema et al., 2014). Nevertheless, these approaches face challenges such as the difficult and complex preparation of sensors(Abe et al., 2011; Knapp et al., 2017), narrow response range(Knapp et al., 2017), and issues with penetration through eukaryotic membranes(Wurm et al., 2010). Therefore, there is an urgent need for versatile fluorescent reporter systems that are easy to prepare, have a wide response range, high sensitivity, and are suitable for use in eukaryotes.

To address the aforementioned challenges, we here propose a novel strategy combining dual-Base editor mediated *In-situ* genome engineering with Nanobody-regulated biosensor-assisted Droplet-sorting to Enhance R-protein secretion (BINDER) in *P. pastoris* (Fig. 1). BINDER enables searching for hyper-producers within an extensive genetic sequence space, and its target editing feature allows the rapid identification of key mutations conferring phenotypic improvements among hundreds of thousands of mutants. As a proof pf concept, recombinant human serum albumin (rHSA) was chosen as a target product. rHSA is a therapeutic protein with high clinical demand for various medical treatments, including blood volume replacement and drug delivery, making its efficient production highly significant. We successfully employed BINDER to engineer three transcriptional regulators, Mxr1, Yap1 and Hac1, which regulate hundreds of genes. A total of 261 sgRNAs were designed, predicting the generation of a mutant library with a substantial size of 113,632. Two critical SNVs were identified, with the *HAC1*_S224L mutation showing a 1.78-fold improvement in rHSA secretion titer, surpassing the *HAC1* overexpressing strain. Additionally, *HAC1*_S224L demonstrated potential in enhancing the secretion titer of other r-proteins. Transcriptome analysis revealed that this mutation improved secretion capability by regulating the Hsp70 co-chaperone cycle, providing a new understanding of the function of the *HAC1* gene and insights into the secretion mechanism. Subsequent fed-batch cultivation of the engineered strain reached the highest reported rHSA titer of 23.43 g/L, demonstrating its substantial potential for industrial applications.

**Figure 1.**
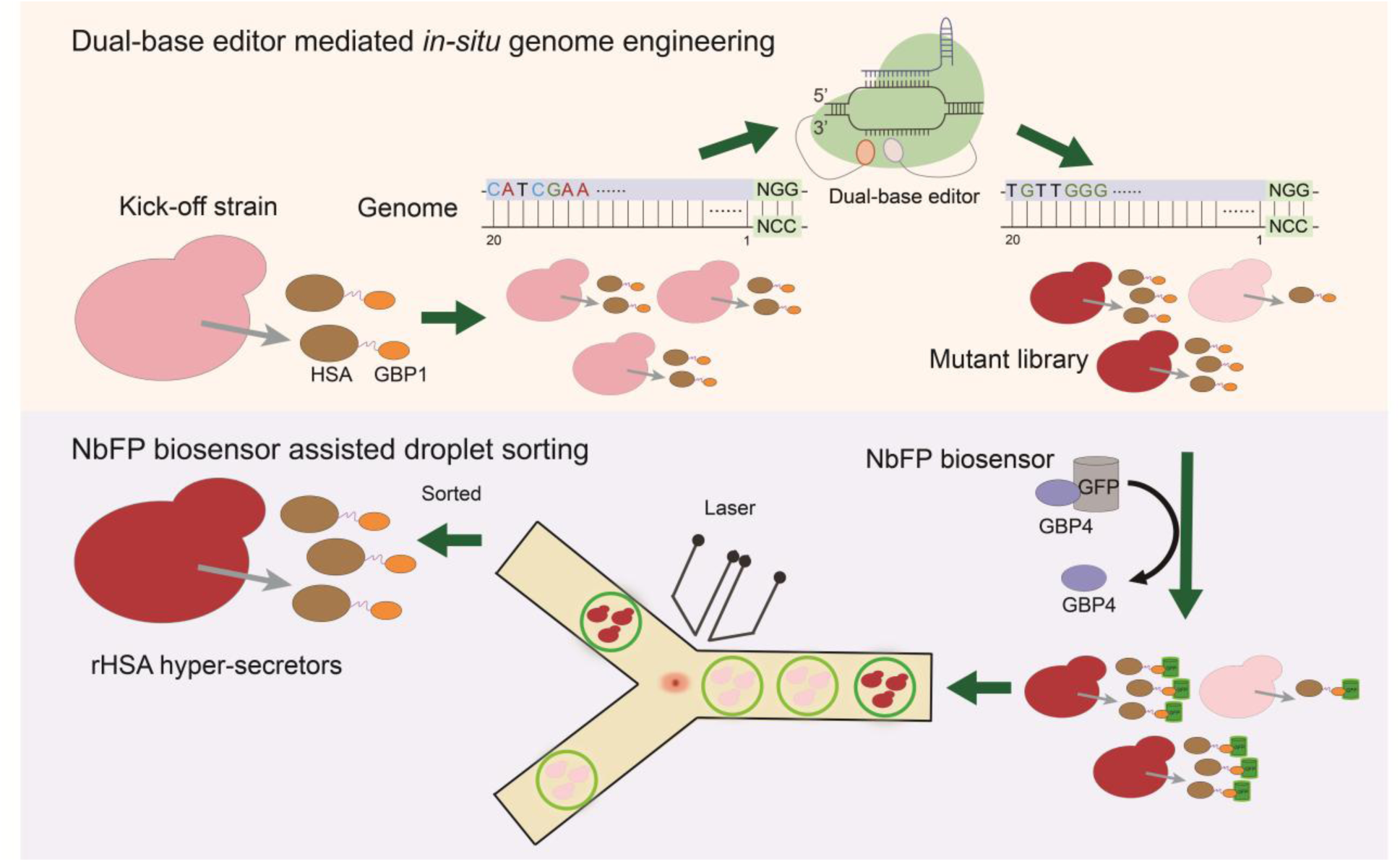
Overview of the BINDER strategy. The dual-base editor was employed to perform *in-situ* engineering and generate a mutant library. The NbFP biosensor-assisted droplet sorting was developed to screen for rHSA hyper-secretors. rHSA: recombinant human serum albumin; NbFP: nanobody-regulated fluorescent protein; GFP: green fluorescent protein; GBP1: GFP-binding protein 1; GBP4: GFP-binding protein 4.

## 2. Results

### 2.1. Establishment and optimization of DBE in *P. pastoris*

We sought to identify an appropriate BE for *in-situ* directed mutagenesis in *P. pastoris*. By employing cytidine or adenine deaminases, cytidines (Cs) or adenines (As) in target regions can be deaminated to uridines (Us) or inosines (Is), respectively. During DNA repair and replication, Us or Is are recognized as thymidines (Ts) or guanosines (Gs) by DNA polymerase, resulting in C>T or A>G conversions. First, we engineered two variants of a cytosine base editor (CBE) to evaluate the feasibility of BE in *P. pastoris*, wherein the cytidine deaminase (CDA) from *Petromyzon marinus* was fused to the C-terminal of dCas9 and nCas9(D10A), thereby generating dCas9-CDA and nCas9-CDA. An RNA polymerase III-based expression cassette of single guide RNA (sgRNA) was integrated into the *HIS4* locus in the genome of the GS115 strain. Subsequently, dCas9-CDA and nCas9-CDA, whose expression were driven by the constitutive promoter *pENO1*, were integrated into the *pENO1* locus of the resultant strain (Fig. 2A). To evaluate mutagenesis efficiency, *ADE2* was selected as a target gene, given that its disruption would result in a discernible pink phenotype. We designed an sgRNA to generate a conversion from a glutamine codon (CAG) to a stop codon (TAG), thereby disrupting the *ADE2* gene. Consequently, colonies exhibiting a deeper pink hue were observed when nCas9-CDA was employed, indicating a higher editing efficiency (Fig. 2B). Next generation sequencing (NGS) results corroborated that nCas9-CDA introduced SNVs more efficiently, with no indels observed (Fig. S1). The performance of nCas9-CDA was further assessed by designing two additional sgRNAs targeting cytosine-enriched regions. Base substitutions were achieved at cytosine positions -14 to -20 upstream of PAM (editing efficiency > 0.5%), where C-to-T conversion was the predominant substitution type (Fig. 2C).

**Figure 2.**
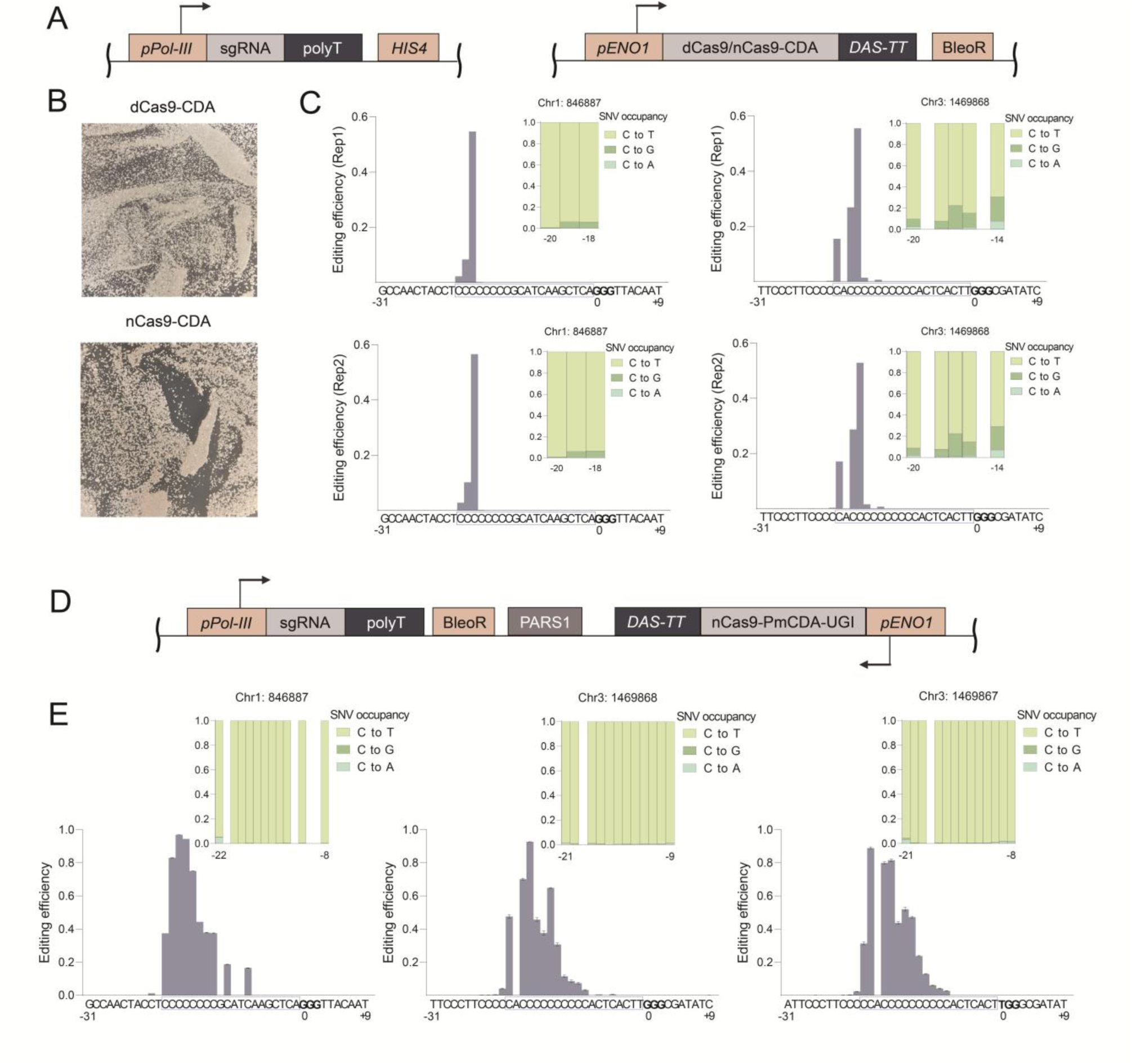
Establishment and optimization of CBE in *P. pastoris*. (A) A schematic view of the integrated CBE. (B) The resulting colonies when *ADE2* gene was targeted. (C) The editing efficiency and mutation types of integrated dCas9-CDA and nCas9-CDA. Mutation types were shown only for the positions with editing efficiency higher than 0.5%. Two biological replicates were performed (Rep1 and Rep2). (D) The episomal version of CBE, nCas9-CDA-UGI. (E) The editing efficiency and base substitution types when episomal nCas9-CDA-UGI was applied. Mutation types are shown only for the positions with editing efficiency higher than 0.5%. Two biological replicates were performed. Error bar represents standard deviation. Chr: chromosome.

When integrating BE into the genome of the host, we observed severe growth inhibition, which may be due to the toxicity and metabolic burden brought by the Cas9 protein variant and deaminase (Fig. S2). Moreover, the long-term expression of BE has been proven to frequently increase off-target effects(Slesarenko et al., 2022). To mitigate the growth toxicity and potential off-target effects, we constructed nCas9-CDA and the sgRNA expression cassette on an episomal plasmid, which could be cured post-editing by passaging in antibiotic-free medium (Fig. 2D). Mutants devoid of BE plasmids could subsequently be subjected to hyper-producer sorting. To further enhance the editing efficiency, an uracil glycosylase inhibitor (UGI) was also fused to the C-terminal of nCas9-CDA (Fig. 2D). The resulting CBE, denoted as nCas9-CDA-UGI, exhibited a significantly elevated mutagenesis efficiency and an editing window extending up to 15-nt (Fig. 2E). To assess the eliminability of BEs, we inoculated and propagated the strains containing the BE plasmid in YPD liquid medium without antibiotics over 24 h. The resulting cultures were diluted and spread onto YPD plates with or without antibiotics for colony counting. The results indicated an elimination rate exceeding 99% (Fig. S3).

Following the construction of episomal CBE, we incorporated adenine deaminases from two recently reported adenine base editors (ABE), namely ABE7.10 and ABE8e, into the N-terminal of nCas9-CDA-UGI. This led to the creation of DBEs, which facilitated a broader mutation spectrum. To evaluate the performance of these DBEs, we examined the editing efficiency at the *ADE2* locus as well as two additional regions that are abundant in adenines and cytosines. Compared to ABE7.10-nCas9-CDA-UGI, ABE8e-nCas9-CDA-UGI demonstrated a higher A-to-G conversion efficiency and an expanded editing window as wide as 15-nt (editing efficiency > 0.5%, Fig. 3).

**Figure 3.**
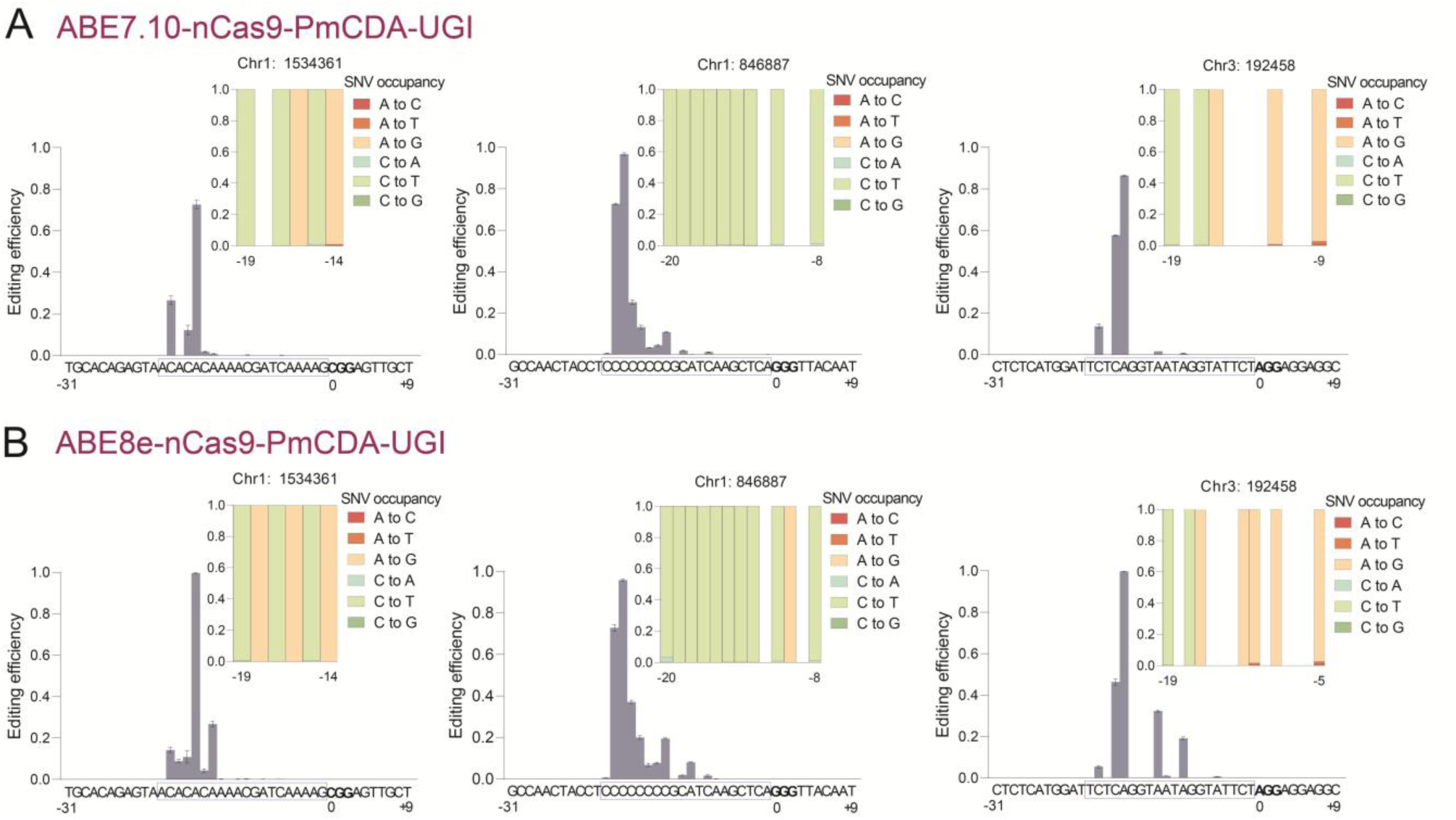
Establishment and optimization of DBE in *P. pastoris*. (A), (B) The editing efficiency and base substitution types when episomal ABE7.10-nCas9-CDA-UGI and ABE8e-nCas9-CDA-UGI was applied, respectively. Mutation types are presented only for positions with an editing efficiency exceeding 0.5%. Two biological replicates were conducted. Error bars represent standard deviation.

### 2.2. Development of a biosensor-assisted droplet sorting platform for r-protein hypersecretion

To develop a scalable high-throughput screening platform for protein hypersecreting strains, we first evaluated a biarsenical compound FlAsH-mediated biosensor, which has been successfully applied to screen r-protein hyper-secreting *Corynebacterium glutamicum*(Yu et al., 2023). This approach tags the secretory proteins with a tetracysteine motif (CCPGCC) and adds FlAsH-EDT_2_ compound to emit green fluorescence. However, our experiments revealed that FlAsH-EDT_2_ penetrated yeast cells within 30 min, engendering intracellular fluorescent signals (Fig. S4). This property disqualified it for use in monitoring r-protein secretion and microfluidic sorting of eukaryotic systems. Consequently, we shifted our focus towards the development of an innovative biosensor.

Nanobodies, antibodies that naturally lack the light chain and are found in the peripheral blood of camels, consist of a single heavy chain variable region (VHH) and two conventional CH2 and CH3 regions. With a molecular weight not exceeding 15 KDa, nanobodies have been shown to be expressed and secreted across a diverse range of systems, including *E. coli*, *C. glutamicum*, yeast, and mammalian cells(de Marco, 2020; Yu et al., 2023). Furthermore, their stability, high specificity, and other advantages have garnered considerable attention in the fields of biotechnology research and medical diagnostics(Muyldermans, 2013). Anti-GFP or eGFP nanobodies (GFP-binding proteins, GBPs) have the ability to modulate the brightness of GFP(Kirchhofer et al., 2010). For instance, the enhanced nanobody GBP1 can specifically augment the fluorescence intensity of GFP, while the inhibitory nanobody GBP4 can specifically attenuate GFP’s fluorescence intensity(Kirchhofer et al., 2010). Based on this principle, we designed a nanobody-regulated fluorescent protein biosensor. Specifically, GBP1 was fused to the target protein, and a peptide linker (termed L10) was added to minimize its interference with the expression of the target protein. GFP or eGFP and GBP4 were independently expressed and purified into an equimolar mixture, which was then added to the system containing GBP1-tagged target proteins. As GBP1 and GBP4 competitively bound to wtGFP or eGFP, GBP1 ultimately displaced GBP4, leading to the regeneration of fluorescent signals from wtGFP or eGFP. This allows for the accurate quantitative determination of the secreted titer of the target r-protein (Fig. 4A).

**Figure 4.**
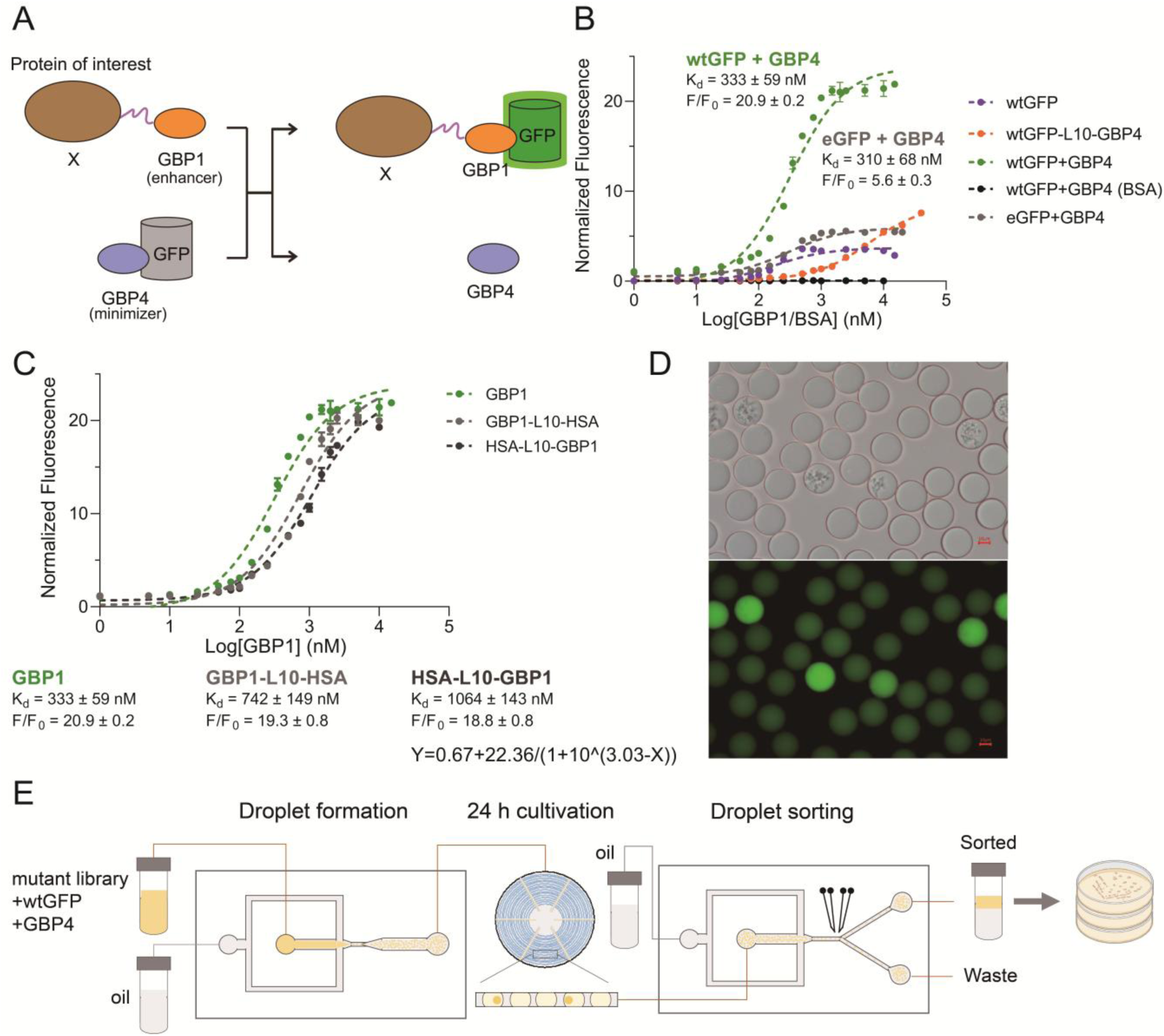
Nanobody-regulated biosensor for monitoring r-protein secretion. (A) Working principle of the nanobody-regulated fluorescent protein biosensor. (B) Optimization of the fluorescent reporter systems. (C) Dose-response curve of GBP1, GBP1-L10-HSA, and HSA-L10-GBP1 as target r-proteins using wtGFP+GBP4 as a probe. The fitted dose-response curve of HSA-L10-GBP1 is shown, with X representing Log_10_[GBP1] and Y representing normalized fluorescence. (D) Microdroplets after 24 h cultivation. The diameter of the droplets was approximately 30 *μ*m, and around 10% of the total droplets contained cells. (E) Overview of the biosensor-assisted high-throughput screening procedure.

When employing wtGFP, wtGFP-L10-GBP4, wtGFP+GBP4, and eGFP+GBP4 as fluorescent reporter systems, GBP1 elicited discernible response signals (Fig. 4B). However, the dynamic ranges of wtGFP and wtGFP-L10-GBP4 were suboptimal. When wtGFP+GBP4 was utilized as a fluorescent reporter system, an equimolar quantity of GBP4 could stably suppress wtGFP fluorescence by over 80% within 5 min (Fig. S5A). A conspicuous fluorescence response was observed at a GBP1 concentration of 10-1000 nM, and the signal remained stable post a 5-min reaction (Fig. S5B). The dynamic range ΔF/F0 attained 21.9±0.2, and the dissociation constant Kd was 323±59 nM. Simultaneously, there was no fluorescence response signal when bovine serum albumin (BSA) was added, indicating a robust specificity to the target protein (Fig. 4B). When eGFP+GBP4 was employed as a fluorescent reporter system, although the dissociation constant Kd reached a comparable level with wtGFP+GBP4 (310±68nM), the dynamic range ΔF/F0 was merely 4.5±0.1. Hence, an equimolar mixture of wtGFP and GBP4 was selected as a fluorescent reporter system for detecting the extracellular titer of r-proteins. The fusion of GBP1 at the N-terminus or C-terminus of HSA can accurately and quantitatively detect the intracellular concentration of HSA, with a detectable concentration between 10-1000 nM, and dynamic ranges ΔF/F0 of 19.3±0.8 and 18.8±0.8, respectively (Fig. 4C). Thus, we developed a high-performance biosensor monitoring r-protein secretion with strong specificity and a wide dynamic range.

To match the diversity of mutants generated by DBE, we integrated this sensor with droplet microfluidic technology to establish a biosensor-assisted high-throughput single-cell sorting platform. Initially, we co-cultured low-producing (termed G1 strain) and high-producing HSA-L10-GBP1 (G1 strain co-expressing *HAC1* gene) strains with probes in deep-well plates, determining that a 2 *μ*M wtGFP+GBP4 mixture was the optimal addition concentration (Fig. S6). We then mixed the pre-cultured high-producing and low- producing strains at a 1:1 ratio, and added the wtGFP+GBP4 probe at the optimal concentration. The resulting mixture was then subjected to the droplet microfluidic cell sorter DREM cell to produce single-cell microdroplets at a speed of 3000/s. After 24 h incubation at 30℃, microdroplets with varying fluorescence intensities could be clearly observed (Fig. 4D, 4E). The droplets with the highest fluorescence signals were then sorted at a speed of 200/s, collected, and spread on YPD plates for 3-day culture. Colonies were randomly picked and PCR was performed to identify high-producing strains by amplifying the Hac1 co-expression cassette. After two rounds of sorting, the proportion of high-producing strains exceeded 95%, indicating the successful establishment of an efficient biosensor-assisted FADS procedure (Fig. S7). This provided a solution for the engineering of efficient secretion cell factories for r-proteins.

### 2.3. *In-situ* transcriptional regulator engineering for high-level secretion of rHSA in *P. pastoris*

The secretion of r-proteins imposes a heightened metabolic burden, as protein synthesis and secretion necessitate an increase in nutrient and energy demands on the host organism. The r-proteins undergo post-translational modifications such as disulfide bond formation, glycosylation, and phosphorylation in the endoplasmic reticulum (ER) during the secretion process. The folding of secreted proteins induces oxidative stress, as disulfide bond formation and ER stress lead to an imbalance in the redox coenzyme regeneration process(Tomàs-Gamisans et al., 2020). Overexpression of heterologous proteins may also lead to the accumulation of a large number of unfolded or misfolded peptides in the ER, activating the unfolded protein response (UPR) pathway. This, in turn, regulates molecular chaperones, folding enzymes, and ER-related protein degradation pathways. Therefore, three transcription factors, namely Hac1, Yap1, and Mxr1, were selected as targets for *in situ* evolution using ABE8e-nCas9-CDA-UGI to improve rHSA secretion in this section.

Mxr1 (methanol expression regulator 1) is an essential transcription factor in the methanol utilization pathway of *P. pastoris*. Evolving Mxr1 has the potential to regulate nutrient utilization efficiency when methanol is used as a carbon source for fermentation. Hac1 is a UPR-regulating transcription factor that can activate the expression of response elements in the UPR pathway, including protein disulfide isomerase (Pdi1), ER chaperone (Kar2), ER oxidoreductase (Ero1), among others. In yeast, the overexpression or regulation of *HAC1* and UPR response genes has been reported to increase the secretion of heterologous proteins(Guerfal et al., 2010; Lin et al., 2023). The oxidative stress response transcription factor Yap1 is required to alleviate the accumulation of reactive oxygen species (ROS) generated during heterologous protein folding, and its overexpression has been shown to improve trypsin secretion in *P. pastoris*(Delic et al., 2014).

A total of 261 sgRNAs were designed, resulting in a mutant library of 113,632 variants, calculated with an editing window of 15-nt (Table S1). The mutant library was subsequently subjected to a single round of sorting to enrich hyper-secreting members. To ensure comprehensive coverage, colonies transformed by electroporation were obtained in a quantity 10-fold the size of the sgRNA library prior to sorting. Post-sorting strains exhibited a higher fluorescence intensity compared to the pre-sorting library, indicating a higher secretion of HSA-L10-GBP1 (Fig. 5A). A total of 188 post-sorting colonies were randomly picked to validate fermentation performance in deep-well plates. Approximately 75% of these strains exhibited a fluorescence intensity higher than the control strain G1 (Fig. 5B). Sanger sequencing of the top 30 strains revealed 10 distinct mutations, eight of which were located at the *HAC1* gene (Table S2). Fermentation was conducted in flasks to further confirm the performance of the 10 mutants (Fig. 5C, Fig. S8). With the exception of M5, the sensor detection results (fluorescence intensity fold change) of the mutants were consistent with the SDS-PAGE results (titer fold change). The relative fluorescence intensity of M5 is 2.12 times that of G1, but the titer calculated by SDS-PAGE and imageJ(Schneider et al., 2012) was only 63.19% of G1. The protein bands of SDS-PAGE suggested that the reason for this phenomenon is the severe degradation of HSA-L10-GBP1 in this strain, resulting in the secretion of most proteins with incomplete structures into the fermentation broth. Strains M2, M3, M4, M8, M9 and M10 displayed a significantly improved titer (*P*<0.05), with an increase of up to 1.89-fold.

**Figure 5.**
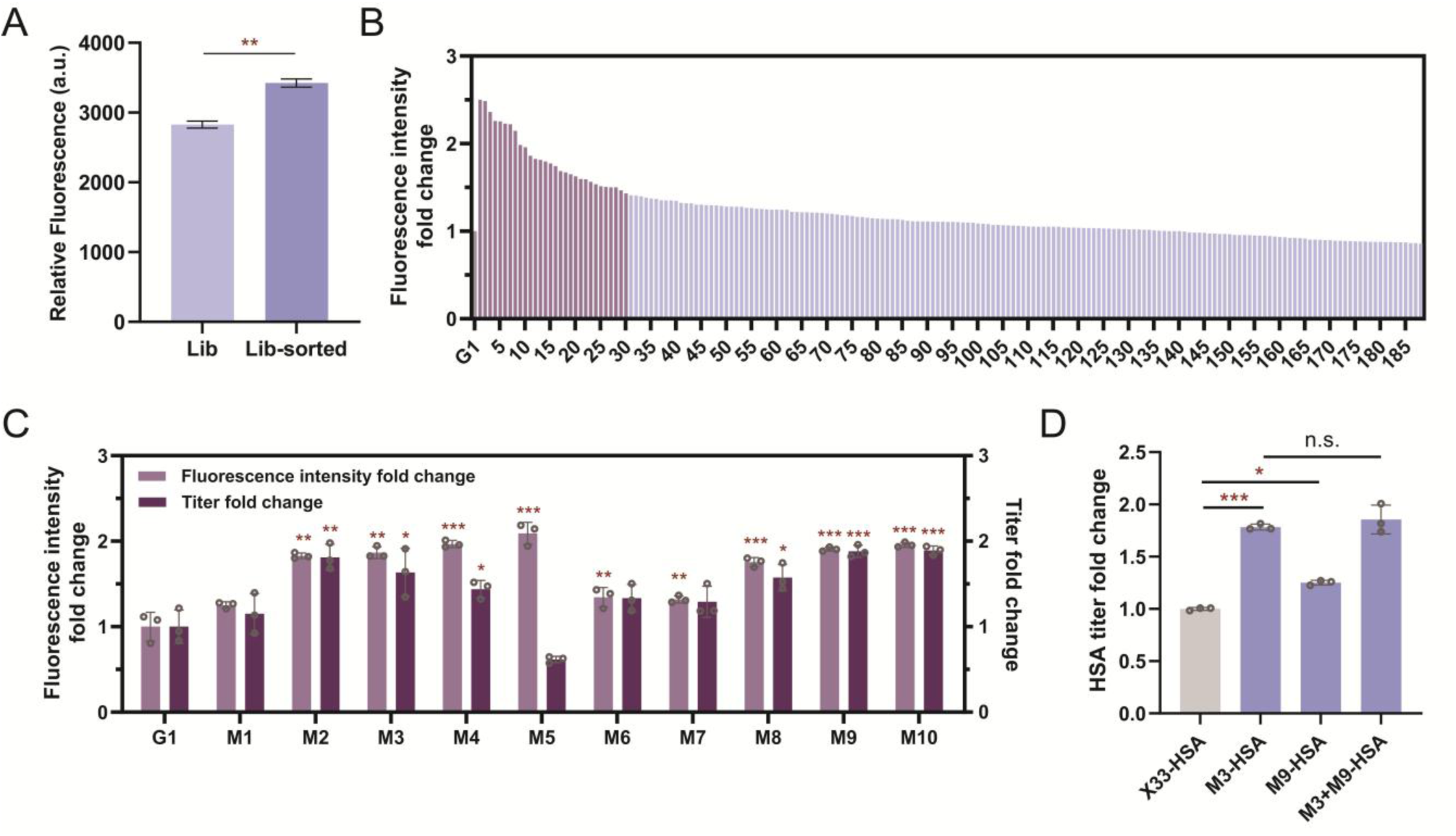
NbFP-biosensor-assisted FADS of the mutant library. (A) R-protein secretion ability of the mutant library before and after FADS. The biosensor was used to detect the fluorescence intensity of the fermentation supernatant after 72 h methanol-induction in flasks containing BMMY medium. Lib and Lib-sorted refer to the mutant library before and after sorting, respectively. Two biological replicates were performed. (B) Performance validation of the colonies post-sorting. The biosensor was used to detect the fluorescence intensity of the fermentation supernatant after 48 h methanol-induction in 48 deep-well plates containing BMMY medium. The 30 strains with the highest fluorescence intensity (shown in purple) were picked to identify genotype by Sanger sequencing. (C) Performance validation of the 10 mutants in flasks. After 72 h methanol-induction in flasks containing BMMY medium, fermentation supernatant was taken to detect fluorescence intensity using the biosensor. SDS-PAGE was performed to isolate HSA-L10-GBP1 and grayscale was analyzed using imageJ to calculate the titer of secreted protein. Three biological replicates were performed. (D) HSA production of the strains with M3, M9, and M3+M9. SDS-PAGE and imageJ were used to quantify HSA concentration after 72 h methanol-induction in flasks containing BMMY medium. Three biological replicates were performed. n.s.: not significant. \*\*\**P*<0.001, \*\**P*<0.01, \**P*<0.05.

To verify that these mutations were instrumental in enhancing the titer, the six mutations were reintroduced into the G1 strain by using CRISPR-Cas9 mediated recombineering. Mutation M3 (*HAC1*_S224L) and M9 (*YAP1*_A44T) increased the protein titer by 1.65-fold and 1.13-fold, respectively, while the remaining mutants demonstrated a titer not higher than G1 (Fig. S9). The underperformance of certain mutants could potentially be attributed to the off-target effects of BEs. Given that GBP1 was used as a tag for microfluidics sorting, we then reconstructed mutation M3 and M9 into an X33 strain harboring HSA expression cassette (X33-HSA), resulting in the strain M3-HSA and M9-HSA, respectively. After 72 h of fermentation in flasks containing BMMY, the M3-HSA and M9-HSA strains exhibited HSA titers 1.78-fold and 1.24-fold higher than X33-HSA, respectively (Fig. 5D, Fig. S10). A strain carrying both mutations was also constructed; however, no significant improvement in the rHSA titer was observed.

Given that mutation M3 is extremely effective in increasing the secretion of HSA-L10-GBP1 and HSA, we further explored the potential of M3 in improving the secretion of other r-proteins. Two new proteins were constructed for testing purposes. HSA has a half-life of up to 3 weeks in plasma and is thus widely applied in drug delivery. Fusion of drug proteins with HSA can effectively improve pharmacokinetics and extend the circulating half-life of the drug(Shen et al., 2021). In 2022, Linqi Zhang’s group reported a new nanobody 3-2A2-4, which is a broadly neutralizing and protective nanobody against SARS-CoV-2(Li et al., 2022b). We then constructed the first testing protein by fusing 3-2A2-4 to the C-terminal of HSA. Another protein chosen for testing was phytase, which can hydrolyze phytate and release phosphate groups, and is widely used as a feed supplement to improve the utilization of phosphorus by animals. We introduced the M3 mutation into strains secreting the HSA-3-2A2-4 fusion protein and phytase, and the titers of the two proteins increased by 1.38-fold and 1.12-fold, respectively, after 72 h of fermentation in flasks (Fig. S11). These results suggest that the *HAC1*_S224L mutation has potential in promoting the secretion of other r-proteins.

### 2.4. Transcriptome analysis of M3-HSA strain

Given that Hac1 functions as a regulator, transcriptome analysis of X33-HSA and M3-HSA strains was conducted to analyze changes in global gene expression. Compared to strain X33-HSA, 289 and 261 genes showed significant differences in expression levels in M3-HSA (*FDR*<0.05, Log_2_Foldchange>1.2, Table S3), several of which were related to protein folding or dependent on Hac1 (Table S4). Most of these genes were involved in the Hsp70 co-chaperone cycle, which plays a pivotal role in protein quality control(Kohler and Andréasson, 2020). In this cycle, Hsp40 and nucleotide exchange factor (NEF) serve as essential co-chaperones, regulating and enhancing the activity of Hsp70 at different stages. Hsp40 enhances substrate binding by promoting ATP hydrolysis and bridging unfolded protein to Hsp70, while NEF controls the release of protein by accelerating ADP to ATP exchange (Fig. 6A). Through this highly regulated cycle, unfolded, partially folded, and misfolded proteins are directed to their respective downstream processes by different co-chaperones in the Hsp70 system for folding or degradation.

**Figure 6.**
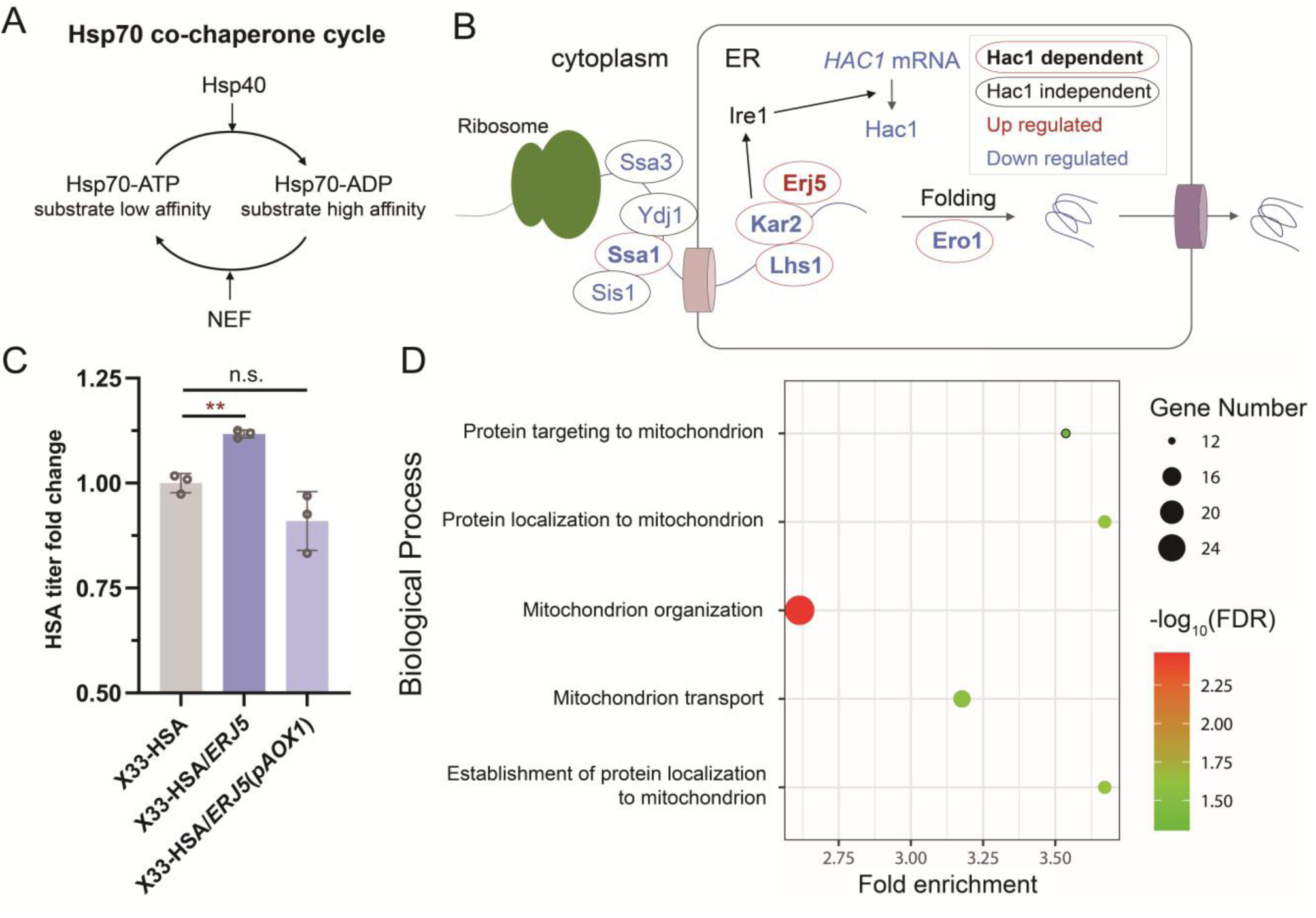
Hsp70 co-chaperone cycle. (A) Hsp70 co-chaperone cycle nucleotide. NEF, exchange factor. (B) Mechanism illustration of the improved secretion ability of the M3 mutant. Proteins encoded by genes with up- and down-regulated transcription levels are shown in red and blue, respectively. Proteins encoded by genes dependent on Hac1 are shown in bold and circled in red. Proteins encoded by genes independent of Hac1 are shown in regular font and circled in black. (C) HSA titer fold change of the strains co-expressing the *ERJ5* gene. X33-HSA/*ERJ5*: X33-HSA strain co-expressing *ERJ5* using its native promoter. X33-HSA/*ERJ5*(*pAOX1*): X33-HSA strain co-expressing *ERJ5* using the promoter *pAOX1*. Three biological replicates were performed. n.s.: not significant. (D) Bubble chart of GO enrichment analysis results. Biological processes with *FDR*<0.05 are shown.

In this work, *S.cerevisiae* MFα-leader sequence was used as a signal peptide to secrete r-proteins in a post-translational translocation manner. The nascent polypeptides are maintained in an unfolded state by Hsp70 co-chaperone cycle in cytoplasm. Upon translocation into the ER, the Hsp70 co-chaperone cycle in ER mediates further processing (Fig. 6B). Most of the genes in this process, including *HAC1* itself, were down-regulated, with the exception of *ERJ5*. Since Hac1 can upregulate its own transcription to alleviate ER stress(Guerfal et al., 2010), we speculate that the down-regulation of the *HAC1*_S224L gene was probably due to the mitigation of ER stress, suggesting it may be a more efficient mutant than the wild-type *HAC1*. In yeast, the gene transcription levels of *SSA1*, *KAR2*, *LHS1*, *ERO1* and *ERJ5* are positively correlated with the transcription levels of wild-type *HAC1* and thus have been identified as *HAC1*-dependent genes(Pincus et al., 2014). Therefore, the decreased transcription levels of *SSA1*, *KAR2*, *LHS1*, and *ERO1* in M3 may be related to the decreased expression level of *HAC1*, while the increased expression level of *ERJ5* may suggest changes in the regulatory mechanism of this *HAC1* mutant. We subsequently overexpressed *ERJ5* in the X33-HSA strain using both its native promoter and the strong promoter *pAOX1*. After 72 h fermentation in flasks, the strains co-expressing *ERJ5* using its native promoter showed an 11% improved titer compared to X33-HSA, while the strains co-expressing *ERJ5* using *pAOX1* showed a decreased titer (Fig. 6C). This result suggested that the increased expression level of *ERJ5* is one of the key factors for the hyperproduction phenotype of M3-HSA. However, the expression level of *ERJ5* should be precisely tuned. A high expression level may lead to an imbalance in the Hsp70 co-chaperone cycle, which is not conducive to the secretion of r-proteins.

Gene Ontology enrichment analysis was also performed, revealing that the differentially expressed genes were significantly enriched in the biological processes related to mitochondrion, with most of these genes being up-regulated (Fig. 6D, S12). We speculate that the M3-HSA strain upregulated these pathways to provide energy for the synthesis of r-protein.

### 2.5. rHSA production by fed-batch fermentation

Given that the co-expression of *HAC1* gene is widely applied to improve r-protein secretion in yeast, we overexpressed the wild-type *HAC1* gene using the *pAOX1* promoter in X33-HSA, resulting in the strain X33-HSA/*HAC1*. Consequently, M3-HSA exhibited a higher secretion titer of HSA compared to X33-HSA/*HAC1*, highlighting the great potential of *in-situ* engineering (Fig. S13). Since the fermentation and transcriptome analysis results indicated that *HAC1*_S224L was a more effective *HAC1* mutant, *HAC1*_S224L gene was subsequently overexpressed in M3-HSA strain using the *pAOX1*. The resulting strain, M3-HSA/*HAC1*_S224L, showed a 2.26-fold higher titer than X33-HSA after 72 h fermentation in flasks (Fig. 7A). To validate that the improved secretion of HSA is scalable, fed-batch cultivations were performed in a 5-L bioreactor (Fig. 7B). A glycerol-methanol DO-stat feeding strategy was employed using basal salt medium (BSM) with half the salt concentration. In the initial phase, glycerol served as sole carbon and energy source. After approximately 24 h of cultivation, methanol was fed as sole carbon source and served as an inducer of rHSA expression for 192 h. The biomass formulation during fed-batch cultivation is shown in Fig. S14. The rHSA titer of M3-HSA/*HAC1*_S224L reached 23.43 g/L after 192 h of induction, which was 2.75-fold higher than that of the X33-HSA strain, representing the highest titer reported so far (Fig. 7C, S15). The HSA yield of M3-HSA/*HAC1*_S224L was 0.198 g HSA/g DCW, representing a 2.42-fold increase compared to X33-HSA (Fig. S16). Taken together, these results underscore the reliable and stable secretion capability of M3-HSA/*HAC1*_S224L.

**Figure 7.**
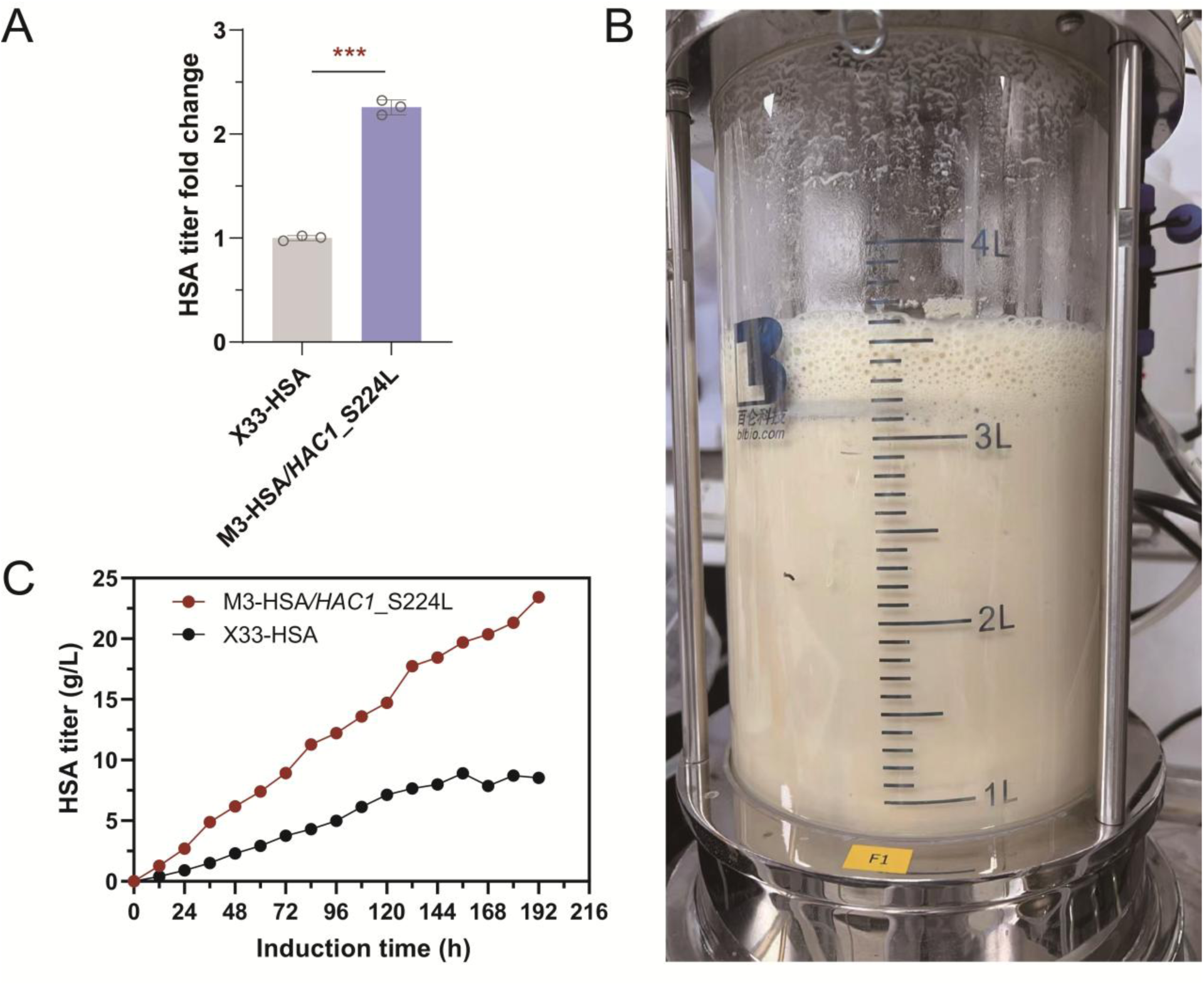
HSA production of M3-HSA/*HAC1*_S224L and X33-HSA. (A) HSA production of M3-HSA/*HAC1*_S224L and X33-HSA. SDS-PAGE and imageJ were applied to quantify HSA concentration in culture supernatant after 72 h methanol-induction in flasks containing BMMY medium. Three biological replicates were performed. (B) Bioreactor for rHSA fermentation. (C) HSA titer of engineered strains in fed-batch cultivation. Data from methanol-induction phase are shown.

## 3. Discussion

*In-situ* genome engineering circumvents the effects of altered gene dosage or genetic context, thereby providing new insights into endogenous genetic networks, and developing superior cell factories. Evolutionary strategies such as adaptive laboratory evolution and random mutagenesis are widely used for *in-situ* engineering of cell factories. However, the presence of genetic hitchhikers complicates the identification of critical mutations. Targeted engineering, which focuses mutations on genes highly correlated with desired phenotypes, holds great potential to accelerate the process of evolution and beneficial mutation identification. Nevertheless, the limited genetic tools in *P. pastoris* have restricted high-throughput genomic editing studies to targeted gene knockouts, leaving the potential of single-base perturbation underexplored. Additionally, when improving the expression and secretion efficiency of r-proteins in hosts through large-scale mutagenesis, a major bottleneck is the lack of high-throughput methods for detecting extracellular protein titers, which hinders the rapid isolation of hyper-secretors. Consequently, high-throughput engineering of r-protein cell factories is mainly limited to the modification of specific industrial enzymes and antibody producers.

To bridge this gap, we propose BINDER, which combines DBE-mediated *in-situ* targeted mutagenesis and NbFP biosensor-assisted droplet sorting. BINDER facilitates the efficient generation of a mutation library, producing a substantial library of 113,632 mutants using 261 sgRNAs. The NbFP biosensor-assisted droplet sorting achieves a generation rate of 3,000 droplets per second and a sorting rate of 200 droplets per second, aligning with the throughput of mutant creation. In contrast, most high-throughput screening studies in *P. pastoris* have only created and evaluated tens of thousands of mutants with gene disruption, limiting the scope of investigation(Ito et al., 2022; Tafrishi et al., 2024). By employing BINDER, we successfully obtained two rHSA hyper-secretors and identified key SNVs conferring up to 1.78-fold enhanced rHSA titer from hundreds of thousands of variants within two months. Typically, a much longer duration is required to isolate hyper-producers from random mutated libraries (e.g., adaptive laboratory evolution necessitates 100-1000 generations of passaging(Wu et al., 2022)). Moreover, strains obtained through adaptive laboratory evolution or random mutagenesis often harbor hundreds of mutations(Huang et al., 2015; Sun et al., 2023), making it laborious to pinpoint the critical ones. Therefore, BINDER offers significant advantages in high-performance strain acquisition and the efficient identification of key mutation sites. Given the high transferability of BEs and the NbFP biosensor’s independence from the properties of the target protein, BINDER might be expanded to a variety of microbial species and r-proteins.

Significant advancements have been made in improving stress tolerance and product titers by introducing SNVs into critical genes(Alper et al., 2006; Alper and Stephanopoulos, 2007; Deng et al., 2024; Garst et al., 2017; Jakočiūnas et al., 2018; Wang et al., 2009; Yuan et al., 2024). Moreover, numerous studies have elucidated the crucial role of SNVs and their mechanisms in phenotype control(Liu et al., 2020; Utrilla et al., 2016; Xiao et al., 2017). A particularly valuable discovery in this work is the *HAC1*_S224L mutant, which enhances the secretion capacity for various proteins. Notably, overexpressing the *HAC1* gene is one of the most effective methods for enhancing r-protein secretion titer in *P. pastoris*. The M3-HSA strain exhibited a greater improvement in rHSA secretion titer compared to the X33-HSA/*HAC1* strain, highlighting the significant potential of introducing SNVs in *HAC1*. Transcriptome sequencing results suggest that the improved secretion may be attributed to the regulated Hsp70 co-chaperone cycle. The ultra-high expression of *ERJ5* has been shown to be detrimental to the secretion of r-proteins in *P. pastoris*(Zahrl et al., 2022), consistent with our findings. However, the discovery of the *HAC1*_S224L mutation indicates that fine-tuning *ERJ5* expression can enhance secretion capacity, offering new insights into the secretion mechanism. Further research is warranted to deeply explore the mechanism by which *HAC1*_S224L enhances secretion. Additionally, by overexpressing the *HAC1*_S224L gene mutant, the resultant strain M3-HSA/*HAC1*_S224L exhibited the highest reported rHSA titer in fed-batch cultivation, underscoring the industrial applicability of this mutation.

During the preparation of this manuscript, Wu et al. reported the first establishment of CBE in *P. pastoris*(Wu et al., 2024). While we did not evaluate the multi-site editing capabilities of BE as they did, nor did we use additional Cas protein variants to broaden the PAM, we constructed and tested the first DBE in *P. pastoris* to further diversify mutation types and focused on exploring its application potential. Future studies could similarly evaluate and optimize DBE. A major drawback of BE-mediated mutagenesis is off-target editing, which reduces the efficiency of identifying key mutations. To minimize off-target effects, deaminases with reduced off-target activity could be used to construct BEs. We tested a cytosine deaminase variant lacking the single-strand DNA binding domain(Li et al., 2022a) and successfully observed C-to-T conversions. However, no mutations were observed when fused with adenine deaminase (data not shown), suggesting that the fusion pattern and editing activity of deaminases require further optimization. Another limitation is the restricted mutation spectrum of deaminases. Recently, DNA glycosylases have been employed to develop BEs with a broader range of mutation types(Wang et al., 2022; Ye et al., 2024; Zhao et al., 2021). Combined with the construction of multiple gRNA arrays, *in-situ* mutations with greater diversity could be introduced combinatorially at multiple genomic loci in *P. pastoris*. The curable DBE developed here also allows for iterative mutagenesis for continuous phenotype optimization. Overall, we are confident that our strategy holds significant potential for optimizing the performance of microbial cell factories.

## 4. Method

### Basic strains and r-proteins

The strains used in this study are based on the *P. pastoris* GS115 and X33. The pPICKαAm vectors containing the expression cassettes of r-protein HSA-L10-GBP1, HSA, HSA-3-2A2-4 or phytase were linearized with *Nru*Ⅰ and integrated into the T38473 locus(Ito et al., 2018) of *P. pastoris* X33 genome. Expression of the heterologous genes was controlled by the methanol inducible alcohol oxidase promoter (*pAOX1*) and terminator (*AOX1tt*). *S.cerevisiae* MFα-leader sequence was fused with each gene encoding r-protein for secretion. The transformed host strains were spread onto YPD plates (10 g/L yeast extract, 20 g/L peptone, 20 g/L glucose, 2% agar) with 100 mg/L G418. PCR was performed and Sanger sequencing was applied to confirm the successful integration of vector in clones. The protocols of the preparation of competent cells and electroporation were described in the following text.

### Plasmids construction

All plasmids used in this study are listed in Table S4. Key primers and oligonucleotides used in the study are listed in Table S5. The *E.coli* DH5α strains were used for plasmid propagation. For most of the plasmids, DNA fragments were amplified by PCR (KOD-FX polymerase, TAKARA Bio, Japan) and assembled into vectors using Gibson assembly (Uniclone One Step Seamless Cloning Kit, Beijing Genesand Biotech Co.,Ltd, China). For episomal plasmids responsible for base editing or CRISPR-Cas9 mediated recombineering, vectors containing a *ccdB* gene expression cassette with two *Lgu*I sites in opposite orientations were initially constructed via Gibson assembly, resulting in pCas9-tRNA1-*ccdB*-sgRNA-Zeo-Epi, pABE7.10-nCas9-CDA-*ccdB*-sgRNA-Zeo-Epi, and pABE8e-nCas9-CDA-*ccdB*-sgRNA-Zeo-Epi. Subsequently, a pair of oligonucleotide DNAs was designed as follows: 5’-AGC-(20-nt sgRNA spacer sequence)-3’ and 5’-AAC-(reverse complement of the 20-nt sgRNA spacer sequence)-3’. T4 DNA ligase (New England Biolabs, U.S.A.) was then employed to ligate the annealed product to the *Lgu*I-digested vectors, replacing the *ccdB* expression cassette and yielding plasmids expressing the corresponding sgRNAs. Given that the expression of the *ccdB* gene is lethal in *E. coli* DH5α strains, all transformants carried successfully constructed sgRNA expression plasmids. *E. coli* DB3.1 was used to construct plasmids containing the *ccdB* gene expression cassette. All sgRNAs were designed using an online software (http://chopchop.cbu.uib.no/). Critical DNA sequences are listed in Supporting Information 2. The sequence of the key plasmids used in this study are accessible through the following links:

pnCas9-CDA-UGI-*ccdB*-sgRNA-Zeo-Epi:

https://benchling.com/s/seq-4buPLufRGjJMiNTzA5wJ?m=slm-lp0NhguhZwkXOrNrOCP8

pABE7.10-nCas9-CDA-*ccdB*-sgRNA-Zeo-Epi:

https://benchling.com/s/seq-OyNP13Zn7M6lEkAwhVRO?m=slm-dMJP7SIVclUm9ukuFkwj

pABE8e-nCas9-CDA-*ccdB*-sgRNA-Zeo-Epi:

https://benchling.com/s/seq-jp1MGXQZy0BA75hjrX7D?m=slm-RZFwF4x57UpxdNxBS4Xm

pCas9-tRNA1-*ccdB*-sgRNA-Zeo-Epi:

https://benchling.com/s/seq-oUXqruVPLtt9CzFmSS8w?m=slm-tPMrGkMmdE5QqysNhWdQ

pPICKαAm vector for r-protein secretion:

https://benchling.com/s/seq-K2ZEQxmiU3kTVMq3mgEb?m=slm-Ai10d5TIdm3kyWRExc5G

*E. coli* strains were cultured at 37℃ in Luria-Bertani (LB) liquid medium (1% tryptone, 0.5% yeast extract, 1% NaCl) or on LB-agar plates containing 2% agar. When necessary, the medium was supplemented with kanamycin (50 mg/L) or zeocin (30 mg/L). Specifically, when zeocin was added, the LB medium was prepared with 0.4% NaCl.

### Competent cell preparation and DNA transformation

For the preparation of competent cells, yeast colonies were inoculated into 5 mL of YPD liquid medium and shaken overnight. The cultures were then transferred to 100 mL of YPD liquid medium and shaken at 30°C until the OD_600_ reached 1.5. The cells were collected by centrifugation, resuspended in 100 mL of freshly prepared LDST solution (for 1 L: 100 mM LiAc, 10 mM dithiothreitol, 0.6 M sorbitol, and 10 mM Tris-HCl, pH 7.5), and incubated at 30°C for 30 minutes. The cells were then collected by centrifugation at 4°C and washed three times with ice-cold 1 M sorbitol solution. Finally, the cells were resuspended in 800 μL of sorbitol solution, divided into 80 μL aliquots, and stored at -80°C.

For DNA electroporation, 200 ng of linearized or circular plasmids were mixed with 80 μL of competent cells and added to 2 mm electroporation cuvettes or 25-well electroporation plates. Electroporation was performed using the BTX Harvard apparatus ECM 630 High Throughput Electroporation System with parameters set to 1.5 kV, 200 Ω, and 50 μF. The transformed cells were incubated in 1 M sorbitol solution (1:10 v/v) for 1.5 hours at 30°C. The cells were then collected by centrifugation and spread onto YPD plates containing the appropriate antibiotics. When integrating the vectors into the *HIS4* locus of the GS115 genome, the strains were immediately spread onto MD plates containing 1.34% yeast nitrogen base (YNB), 2% glucose, and 2% agar after electroporation.

### Base editing and evaluation of mutagenesis efficiency

After electroporation of BE plasmids, the strains were spread onto the YPD plates with 100 mg/L zeocin for a 3-day incubation at 30℃. Subsequently, all the colonies in the plates were washed, collected and used as templates to amplify target regions by PCR. NGS was performed to analyze the editing efficiency.

### Elimination of the episomal plasmid

Edited cells were inoculated into YPD liquid media without any antibiotics, and cultured at 30℃, 250 rpm for approximately 24h for plasmid elimination. Subsequently, the culture was diluted and spread onto YPD plates with (termed YPDZ) and without zeocin. Plasmid curing rate was calculated as below:

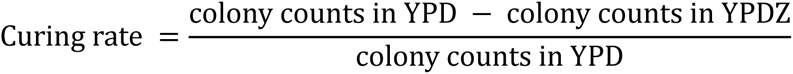

To obtain certain strains without plasmid, colonies were picked and dotted at YPD and YPDZ plates. The colonies failed to grow on YPDZ plates were confirmed as plasmid-free strains.

### Preparation of NbFP biosensor

The genes encoding wtGFP and eGFP were fused with a 6×His tag to facilitate purification using nickel affinity chromatography. *E. coli* BL21(DE3) cells expressing GFP were inoculated into 5 mL of LB medium containing kanamycin and cultured overnight at 30℃ and 220 rpm. The overnight culture was then diluted 1:100 into 100 mL of LB medium with kanamycin and grown until the OD_600_ reached 0.4. Protein expression was induced with IPTG for 24 h. Cells were harvested by centrifugation at 8000×g for 10 minutes at 4℃, washed twice with buffer A (30 mmol/L imidazole, 20 mmol/L sodium phosphate, 500 mmol/L sodium chloride, pH 7.4, filter sterilized, ultrasonicated for 10 min), and resuspended. Cells were lysed using a homogenizer at a pressure of 15,000-20,000 psi. The lysate was then centrifuged at 13,000 rpm for 30 min at 4℃, and the supernatant was filtered through a 0.22 μm membrane and kept on ice.

*P. pastoris* was employed for the secretion of GBP1, GBP4, GFP-GBP4, GBP1-L10-HSA, and HSA-L10-GBP1 proteins. A 6×His tag was fused to these proteins to facilitate their purification during the preparation and characterization of NbFP biosensors. *P. pastoris* clones were inoculated into 5-10 mL of BMGY medium and cultured overnight at 30℃ and 250 rpm. Cells were harvested by centrifugation at 1500×g for 2 min at 30℃, resuspended in BMMY medium, and transferred to flasks containing 10-100 mL of BMMY medium with an initial OD_600_ of 1. Cultures were incubated at 30℃ and 250 rpm for 72 h, with 1% methanol added every 24 h to induce protein expression. The culture supernatant was collected by centrifugation at 10,000 rpm for 10 min at 4℃ and filtered through a 0.22 μm membrane.

Protein purification was performed using an AKTA system at a flow rate of 5 mL/min. The nickel column (Ni-NTA HF) was washed with deionized water and equilibrated with buffer A. The sample was loaded using a syringe, and impurities were washed out with buffer A until the baseline stabilized. Target proteins were eluted with buffer B (500 mmol/L imidazole, 20 mmol/L sodium phosphate, 500 mmol/L sodium chloride, pH 7.4, filter sterilized, ultrasonicated for 10 minutes), and fractions were collected upon observing protein peaks. Collected protein samples were dialyzed against HEPES buffer, and protein concentration was measured using a NanoDrop microvolume UV-Vis spectrophotometer (ThermoFisher Scientific, USA). Samples were aliquoted into 1.5 mL tubes and stored at -80℃.

### Enriching HSA-L10-GBP1 hyper-producers by FADS

The plasmid-free mutant library was pre-cultured in BMGY medium overnight at 30°C. Subsequently, the cells were collected and resuspended in fresh BMMY medium containing 1.5% methanol with an OD_600_ of 0.5 (λ = 0.1), ensuring that the majority of droplets contained no more than one cell(Mazutis et al., 2013). Following a 30-min ultrasound treatment to isolate adherent cells, an equimolar mixture of GBP4 and wtGFP was added to the culture at a final concentration of 2 μM. A standard flow-focusing droplet nozzle was then used to generate 14-pL droplets at a rate of approximately 3,000 per second. Over 30 min, around 5,400,000 droplets were produced, with nearly 9.1% containing single cells, ensuring a 4-fold coverage of the mutant library size. The droplets were collected in Teflon tubes and incubated at 30°C for 24 hours. Droplets exhibiting the highest fluorescence were then sorted at a rate of 200 per second over an 8-h period. Subsequently, 500 μL of fresh YPD medium and 500 μL of demulsifier 1H,1H,2H,2H-Perfluoro-1-octanol (Sigma-Aldrich, USA) were added to the tubes containing the sorted droplets and incubated for 1 min. The post-sorted library in the supernatant was then collected and stored.

### Fermentation in deep-well plates and flasks

Colonies from the post-sorted library were randomly selected and inoculated into BMMY medium containing 1.5% methanol in 48 deep-well plates, with the G1 strain serving as a negative control. After culturing at 30°C and 1000 rpm for 48 hours, the cells were centrifuged, and the HSA-L10-GBP1 titer in the supernatant was determined using NbFP biosensors (2 μM GBP4 + 2 μM wtGFP).

For flask fermentation, the colonies were inoculated in 5 mL of BMGY medium and cultivated overnight at 30°C and 250 rpm. The cells were then collected and transferred to 10 mL of BMMY medium containing 1.5% methanol, with an OD_600_ of 1, and shaken at 30°C and 250 rpm for 72 h in 100 mL flasks. Methanol was supplemented at 1%(v/v) every 24 h. The cells were centrifuged after 72 h induction, and the titer of the r-protein in the supernatant was determined using NbFP biosensors or grayscale analysis with ImageJ.

### Protein quantification

The concentration of r-proteins was determined using sodium dodecyl sulfate polyacrylamide gel electrophoresis (SDS-PAGE) on 8%, 12%, or 15% polyacrylamide gels, depending on their molecular weight. After electrophoresis, the gels were stained with Coomassie Brilliant Blue (CBB) fast staining solution (QIAGEN, China). Photographs of the stained gels were taken with a WD-9413BX image reader (Liuyi Biotechnology, China), and the grayscale of the target bands was quantified using the ImageJ tool. Commercial BSA was used as a standard to generate a standard curve for protein concentration calculation.

### Candidate mutation reconstruction by CRISPR-Cas9 mediated recombineering

To validate whether the mutations created by DBE were critical for improved r-protein secretion, CRISPR-Cas9 mediated recombineering was used to introduce the in-situ mutations into wild-type strains. Specifically, 1 μg of donor DNA containing the target SNVs and 200 ng of the pCas9-tRNA1-*ccdB*-sgRNA-Zeo-Epi plasmid harboring the corresponding sgRNA sequence were electroporated into 80 μL competent cells. After a 3-day incubation at 30℃, transformants on YPDZ plates were randomly picked, and Sanger sequencing was performed to verify their DNA sequences. Colonies with the correct mutations were then cultured in YPD liquid media for plasmid curing. The plasmid-free strains were successfully reconstructed mutants and stored at -80℃.

### Transcriptome analysis

Single colonies were inoculated into BMGY medium and shaken overnight at 30℃ and 250 rpm. The pre-cultures were then transferred into BMMY medium containing 1.5% methanol with an initial OD_600_ of 1. After 12 hours of shaking, 500 μL of culture was taken and centrifuged at 1,500×g for 5 minutes at 4℃. Cells were collected, and total RNA was extracted using the RiboPure™ RNA purification kit (Thermo Fisher Scientific, Cat. No. AM1926). One microgram of RNA was used to prepare the library, and Oligo(dT) beads were used to isolate PolyA tail-containing RNA (mRNA). The mRNA was then fragmented and reverse transcribed to obtain cDNA. The library was constructed using standard methods and sequenced using the Illumina NovaSeq PE150 platform. Cutadapt (v1.9.1) (Martin, 2011) was used to remove adapters and filter low-quality data to obtain clean data. Hisat2 (v2.2.1)(Kim et al., 2015) was then applied to align the sequencing data with the reference genome sequence and gene model annotation files of *P. pastoris* GS115 (https://www.ncbi.nlm.nih.gov/datasets/genome/GCF_000027005.1/). Using the transcriptome data in FASTA format as a reference gene file, HTSeq (v0.6.1)(Putri et al., 2022) was used to calculate gene expression levels, and the DESeq2 Bioconductor software package(Love et al., 2014) was used for differential expression analysis. The threshold for differentially expressed genes was set at FDR ≤ 0.05. GO enrichment analysis was performed using the DAVID tool(Huang da et al., 2009; Sherman et al., 2022).

### Fed-batch fermentation

Clones of the engineered strains were picked and inoculated into 300 mL of BMGY medium in flasks and cultured at 30℃ and 250 rpm until the OD_600_ reached 10. For the fed-batches, 5-L bioreactors filled with 2.3 L of 1/2 BSM-medium [for 1 L: 13.4 mL H_3_PO_4_ (85%), 0.47 g CaSO4, 7.5 g MgSO_4_⋅7H_2_O, 9.1 g K_2_SO_4_, 2.1 g KOH, 40 g glycerol, 0.1 mL Silicone Antifoam (Sigma-aldrich, U.S.A.), 0.96 mg biotin, 4.78 mL PTM1] were used. The pH was maintained at 5.0 using 25% ammonia. The entire pre-culture was inoculated into the bioreactors. The temperature was set to 30℃, and the oxygen saturation was maintained at 30% by adjusting the stirrer and air flow throughout the process.

During the initial batch phase, stirring was set between 300 and 950 rpm, and the air flow was controlled between 3 and 7 L/min. Antifoam was added as needed to control foam. The batch phase typically lasted about 14 h, with the OD_600_ usually reaching 80.

When a sharp increase in oxygen saturation was observed, 50% (v/v) glycerol was fed at 50 mL/h using a DO-stat model. This meant that glycerol was fed only when the oxygen saturation was higher than 30%. After 8 h of supplementation, the OD_600_ exceeded 120. Afterwards, the strains were starved for 40 min, and the pH was adjusted to 5.75 (0 h induction time).

In the following 2 to 3 h, methanol containing 12 mL/L PTM1 and 2.4 mg/L biotin was fed using a DO-stat model at a rate of 10 mL/h. Subsequently, feeding rate was changed to 20 mL/h for 1 h. The feeding rate was then set to 30 mL/h until the fermentation was completed. Samples were taken for analysis every 12 h.

## Author contribution

Y. Y., X. L. and Y. T. contributed equally to this work, including study design, methodology, experimental conduct, and data analysis. Y. Y. drafted the manuscript., S. L. and Kaho Hasegawa participated in the experimental conduct. C. Z., X. X. and Y. W. were responsible for project design and funding acquisition. All authors revised and approved the manuscript.

## Acknowledgment

This work was supported by grants from the National Key Research and Development Program of China (2023YFC3402300), National Natural Science Foundation of China (U2032210), R&D projects in key areas of Guangdong Province (2022B1111050002) and the R&D projects of Hebei Province (22375503D).

## Data Availability Statement

All strains, plasmids are available from the corresponding author by request. DNA sequence of critical plasmids or genes are provided within the paper and Supporting information 2. The data supporting the findings of this work are available within the paper and the Supplementary materials. Data will be made available on request.

